# Press disturbance unveils different community structure, function and assembly of bacterial taxa and functional genes in mesocosm-scale bioreactors

**DOI:** 10.1101/805788

**Authors:** E. Santillan, F. Constancias, S. Wuertz

## Abstract

Sustained perturbations, or press disturbances, are of interest in microbial ecology as they can drive systems to alternative stable states. Here, we tested the effect of a sustained doubling of organic carbon loading on structure, assembly and function of bacterial communities. Two sets of replicate 5-liter sequencing batch reactors were operated at two different organic carbon loading levels (323 and 629 mg COD L^-1^) for a period of 74 days, following 53 days of acclimation after inoculation with sludge from a full-scale treatment plant. Temporal dynamics of community taxonomic and functional gene structure were derived from metagenomics and 16S rRNA gene metabarcoding data. Assembly mechanisms were assessed through a mathematical null model on the effective bacterial turnover expressed as a proportion of total bacterial diversity. Disturbed reactors exhibited different community function, structure and assembly compared to the undisturbed reactors. Bacterial taxa and functional genes showed dissimilar alpha-diversity and community assembly patterns. Deterministic assembly mechanisms were generally stronger in disturbed reactors, associated with common taxa. Stochastic assembly was more important for functional genes and was driven by rare genes. We urge caution when assessing microbial community assembly mechanisms, as results can vary depending on the approach.

## Introduction

Microbes drive all biogeochemical cycles on Earth, with microbial communities providing important ecosystem functions that impact all other forms of life [1]. Community structure, often described in terms of α- and β-diversity, is thought to have an effect on ecosystem function [2]. Yet, our capacity to predict and manage the functions of microbial communities and how they are linked to community structure is still limited [3]. In this regard, engineered systems like sludge bioreactors for wastewater treatment constitute model systems for microbial ecology studies [4], with measurable ecosystem functions such as carbon and ammonia removal which are not only important in practice [5], but also involve complex microbial communities in a controlled environment [6]. In ecology, disturbances are believed to have direct effects on ecosystems by altering community structure and function [7]. Press disturbances that impose a long-term continuous change of species abundances by altering the environment [8] are of interest in microbial ecology as they can drive systems to alternative stable states with different community function and structure [9]. These disturbances could occur in the form of environment modifications that are not directly harmful to organisms, while still providing opportunities to grow for less abundant community members [10]. In sludge bioreactors, a continuous alteration in the substrate feeding scheme can trigger changes in community function and structure, yet whether these changes are reproducible [11] and whether they can be reversed when the disturbance ceases remains unknown.

Community assembly processes are inherently linked with ecosystem function, as they play an important role in shaping community structure [12]. These mechanisms can be either deterministic [13] or stochastic [14], and they may act in combination to drive patterns of community assembly [15-18]. Disturbance is thought to be a main factor driving these underlying mechanisms of community assembly [19], yet a predictive understanding of its effects is missing [20]. Disturbance can promote stochastic assembly mechanisms that lead communities to divergent states of structure and function [21, 22]; therefore, studies assessing its effects require replicated designs [23]. Also, microbial communities within wastewater treatment systems have been shown to harbour a core group of 100-800 abundant (i.e. common) OTUs across plants and countries [24, 25]. Activated sludge community dynamics have been suggested to differ between common and less abundant (i.e. rare) taxa on the grounds of distance-based analyses [26-29], proposing that common OTUs are driven by deterministic mechanisms of assembly. The contribution of the remaining rare taxa to the treatment performance is as yet unknown, but its potential as a seed-bank for gene and taxonomic diversity [30] merits consideration. Indeed, rare taxa and genes could become very abundant under suitable conditions [31] that can be elicited by disturbance.

The contribution of assembly mechanisms is often quantified via null model analyses [32]. In studies of microbial systems, such analyses are usually focused on community data obtained from a fragment of the 16S rRNA gene [17, 18, 33-38], which is known to be highly conserved between different species of prokaryotes [39]. On the other hand, the use of metagenomics would allow inference of assembly mechanisms from the whole community DNA [40], enabling the assessment of variability in genes that are less conserved and that could unveil important aspects of community assembly. Yet, studies usually employ these sequencing methodologies separately as their results are thought to be hard to reconcile [41]. Further, microbial communities can also be assessed in terms of their functional gene structure [42] and while a few studies on microbial community assembly employed GeoChip analysis [22, 43], to date no studies have evaluated assembly mechanisms from functional gene structure via metagenomics. An assessment of community assembly mechanisms combining both 16S rRNA gene amplicon and metagenomics sequencing would provide novel insights towards a better understanding of the effect of disturbance in community assembly, structure and function, while still allowing a comparison with prior findings.

The objective of this work was to test the effect of a press disturbance of doubling organic loading in a replicated set of activated sludge bioreactors. We expected the imposed press disturbance to have an effect in terms of community function, structure and assembly, with the hypothesis of a stronger deterministic effect at the press disturbed level. Samples were analysed using metagenomics, 16S rRNA gene metabarcoding, and effluent chemical characterization. Patterns of α- and β-diversity were employed to assess temporal dynamics of community structure. Assembly mechanisms were evaluated through a mathematical null model on the effective bacterial turnover expressed as a proportion of total bacterial diversity. Finally, the press disturbance was removed during the last 14 days of the study to evaluate if community function, structure and assembly would display signs of recovery.

## Materials and Methods

### Experimental design

Sequencing batch reactors (SBR) with a 5-L working volume were inoculated with activated sludge from a wastewater treatment plant (WWTP) in Singapore and operated on continuous 12-h cycles with intermittent aeration. The experimental setup was detailed elsewhere [44]. Briefly, four reactors were acclimated to laboratory conditions for 53 days while being fed a complex synthetic wastewater containing a variety of carbon and nitrogen compounds. On d54 the biomass of the acclimation reactors was thoroughly mixed and redistributed across seven reactors. From these, three were randomly selected and designated as high organic loading reactors, receiving double the carbon substrate in terms of chemical oxygen demand (COD) in the feed (629 mg COD L^-1^ and 100 mg TKN L^-1^), as a press disturbance for 60 days. The remaining four reactors were operated at low organic loading conditions (323 mg COD L^-1^ and 92 mg TKN L^-1^). During the last two weeks of the study (d114-d127), the feed for the high organic loading reactors was reduced to equal that of low organic loading reactors (details in Table S1). SBR stages were 5 min feed, 200 min anoxic/anaerobic react, 445 min aerobic react, 50 min sludge settle, and 20 min supernatant drain in each cycle. Temperature was controlled at 30°C, pH at 6-9, and dissolved oxygen concentration at 2-6 mg L^-1^ (during the aerobic phase). Two cycles per day accounted for a hydraulic retention time of 24 h. Effluent and influent compositions were measured 2-3 times per week in accordance with Standard Methods [45]. The targets were soluble COD, total alkalinity, and nitrogen species (ammonium, nitrite and nitrate), as well as total Kjeldahl nitrogen (TKN) in the liquid phase using colorimetric tests and ion chromatography. Sludge biomass was measured as total suspended solids (TSS) twice a week, after which biomass wastage was done to target 1500 mg L^-1^ of TSS in order to control the food-to-biomass ratio (F:M) (Table S1). See [44] for detailed information of the source WWTP, inoculum collection, feed preparation, sludge sampling, analytical methods, DNA extractions, and equations for calculation of operational parameters.

### 16S rRNA gene metabarcoding and reads processing

We used primer set 341f/785r, which targets the V3-V4 variable regions of the bacterial 16S rRNA gene [46]. The libraries were sequenced in house on Illumina MiSeq (v.3) with 20% PhiX spike-in, at 300 bp paired-end read-length. Sequenced sample libraries were processed with the *dada2* (v.1.3.3) R-package [47], which allows inference of amplicon sequence variants (ASVs) [48]. Illumina adaptors and PCR primers were trimmed prior to quality filtering. Sequences were truncated after 280 and 255 nucleotides for forward and reverse reads, respectively. After truncation, reads with expected error rates higher than 3 and 5 for forward and reverse reads, respectively, were removed. Reads were merged with a minimum overlap of 20 bp. Chimeric sequences (0.18% on average) were identified and removed. For a total of 104 samples, an average of 19679 reads were kept per sample after processing, representing 49.2% of the average input reads. Taxonomy was assigned using the SILVA database (v.132) [49]. See [44] for further details including rarefaction curves.

### Metagenomics sequencing and reads processing

Libraries were sequenced in house on Illumina HiSeq2500 in rapid mode at a 250 bp paired-end read-length. In total, around 325 million paired-end reads were generated, with 3.4 ± 0.4 million pairedend reads on average per sample (total 48 samples). Illumina adaptors, short reads, low quality reads or reads containing any ambiguous bases were removed using *cutadapt* [50]. High quality reads (91.0 ± 1.4% of the raw reads) were randomly subsampled to an even depth of 4,678,535 for each sample prior to further analysis. Taxonomic assignment of metagenomics reads was done as previously described [51]. High quality reads were aligned against the NCBI non-redundant (NR) protein database (March 2016) using DIAMOND v.0.7.10.59 [52]. The lowest common ancestor approach implemented in MEGAN Community Edition v.6.5.5 [53] was used to assign taxonomy to the NCBI-NR aligned reads. On average, 36.8% of the high-quality reads were assigned to cellular organisms, of which 98.4% were assigned to the bacterial domain. Functional potential data were also obtained from the metagenomics dataset using MEGAN. We employed the four databases available with the rate of assigned/total reads being the highest for the IP2G database (52%), followed by SEED (21%), COG (15%), and KEGG (13%). See [44] for more details including rarefaction curves.

### Bacterial community analysis and statistics

All reported p-values for statistical tests in this study were corrected for multiple comparisons using a False Discovery Rate (FDR) of 5% [54]. Community structure was assessed by a combination of non-metric multidimensional scaling (NMDS) ordination and multivariate tests (PERMANOVA, PERMDISP) on Bray-Curtis dissimilarity matrixes constructed from square-root transformed normalized abundance data using PRIMER (v.7) [55]. Hill diversity indices [56] were employed to quantify α-diversity as described elsewhere [21, 57]. Welch’s ANOVA was used for univariate testing. Bacterial genera, functional genes from the IP2G database at the lowest gene assigned level (metagenomics dataset), as well as ASVs (16S rRNA gene amplicon dataset), were employed for calculating α− and β−diversity metrics, and for quantifying assembly mechanisms.

### Null Model Analyses on Diversity

The effect of underlying assembly mechanisms was assessed through a mathematical null model which assumes that species interactions are not important for community assembly [32] and quantifies the effective bacterial turnover expressed as a proportion of total bacterial diversity. It was developed for woody plants [58] and recently applied to sludge [21] and groundwater [43] microbial communities. The model defines β-diversity as the β-partition 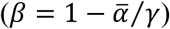, taking into account both composition and relative abundances. To adapt it to handle microbial community data, we considered ‘species’ in the model as ASVs, genera and genes at the lowest IP2G gene level, while each individual count was one read within the corresponding dataset. The model randomizes the location of each individual within the independent replicate reactors for each of the low and high organic loading levels, while maintaining the total quantity of individuals per reactor, the relative abundance of each ‘species’ (i.e. ASV, genus or IP2G gene), and the γ-diversity. We applied it across different time points of the experiment.

Each step of the null model calculates expected mean α-diversities per treatment level and then estimates an expected β-partition. After 10,000 repetitions, the means of the distribution of random β-partitions 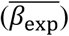 for each treatment level are calculated. The contribution of deterministic assembly mechanisms is then quantified using the deterministic strength (DS) metric, which measures the deviation of the observed beta diversity compared to that expected by chance. DS is equal to the absolute value of the difference between the observed (*β*_*obs*_) and mean expected β-diversity, divided by the observed β-diversity:

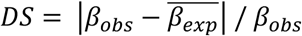. It is the complement of the stochastic intensity (SI) metric defined previously [21]. High values of DS (> 50%) indicate a high deviation of the observed β-diversity from the null β-diversity expectation, thus suggesting a stronger effect of deterministic-based processes. Contrarily, low DS values (< 50%) indicate little difference between observed and null β-diversities, suggesting a more important role of stochastic mechanisms of assembly. Additionally, the expected to observed β-diversity ratio 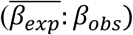 was calculated, for which values below one indicate that the observed β-diversity is greater than expected by chance under the null model, while values above one indicate the opposite.

To further assess community assembly of common and rare fractions, all three datasets were arbitrarily partitioned. Taxa or genes falling into the 99% abundance-rank of accumulated reads of the dataset were classified as common, while the remaining 1% were classified as rare. This was a conservative approach to offset bias when employing relative abundances to quantify taxa or genes.

## Results

### Dynamics of bacterial community structure

Throughout the acclimation phase there was a significant change in bacterial community structure (P-_PERMANOVA_ < 0.005, Table S2) from the starting WWTP inoculum to the acclimated sludge that was later distributed across reactors when the disturbance phase started, both in terms of α-(Fig. 1A) and β-diversity (Fig. 2A).

**Fig. 1.**
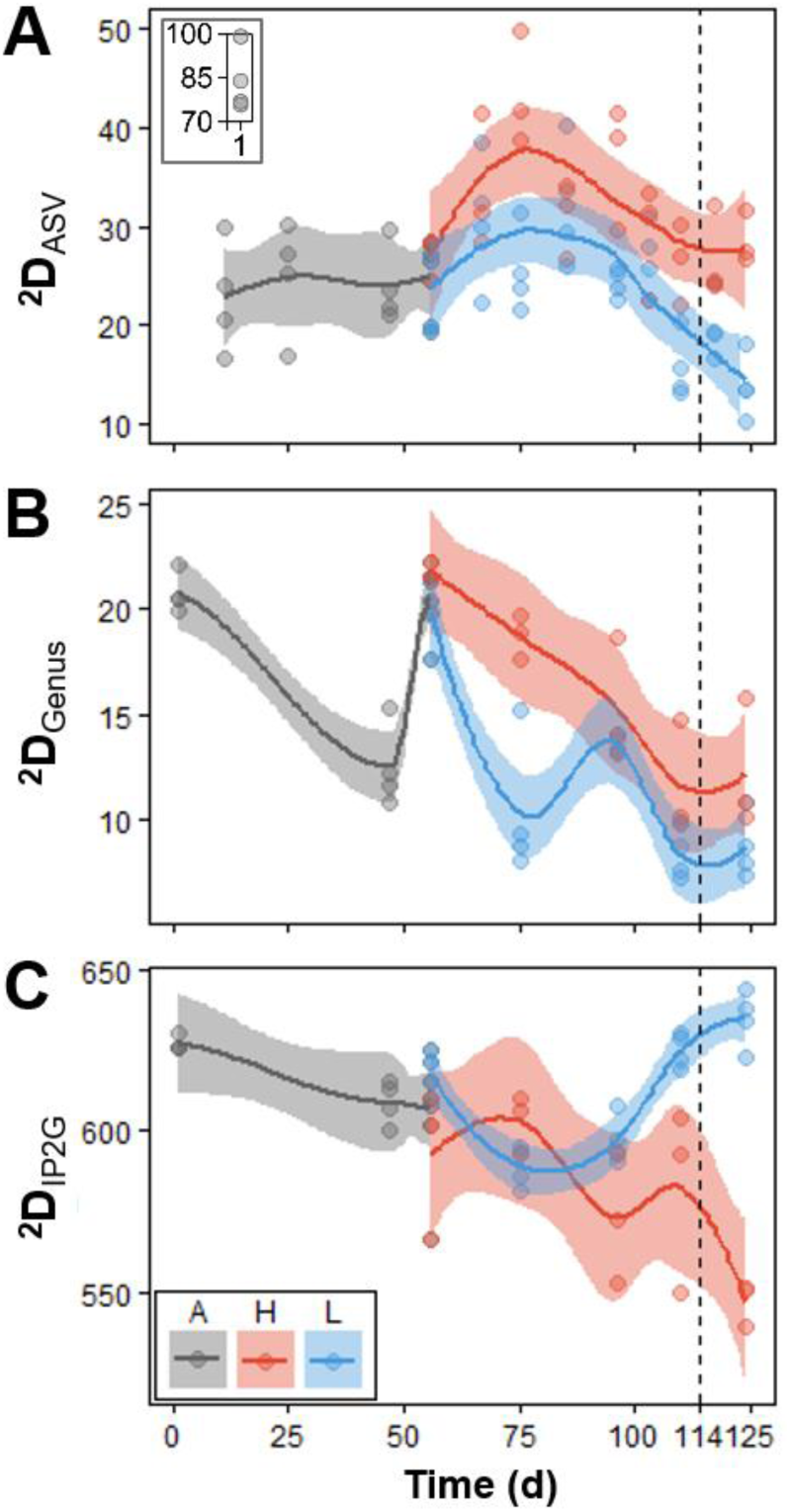
Temporal dynamics of 2^nd^ order true α-diversity (^2^D) for bacterial taxa and functional genes. (**A**) 16S rRNA gene sequencing ASV level, (**B**) metagenomics sequencing genus level, and (**C**) metagenomics sequencing IP2G lowest gene level. Each point represents a different reactor for a given day. Phases: A, acclimation (grey, n = 4); L, low organic loading (blue, n = 4); H, high organic loading (red, n = 3). Vertical dashed line indicates the shift from high to low organic loading. Lines display polynomial regression fitting, while shaded areas represent 95% confidence intervals. Internal panel within (**A**) shows data points at d1.

**Fig. 2.**
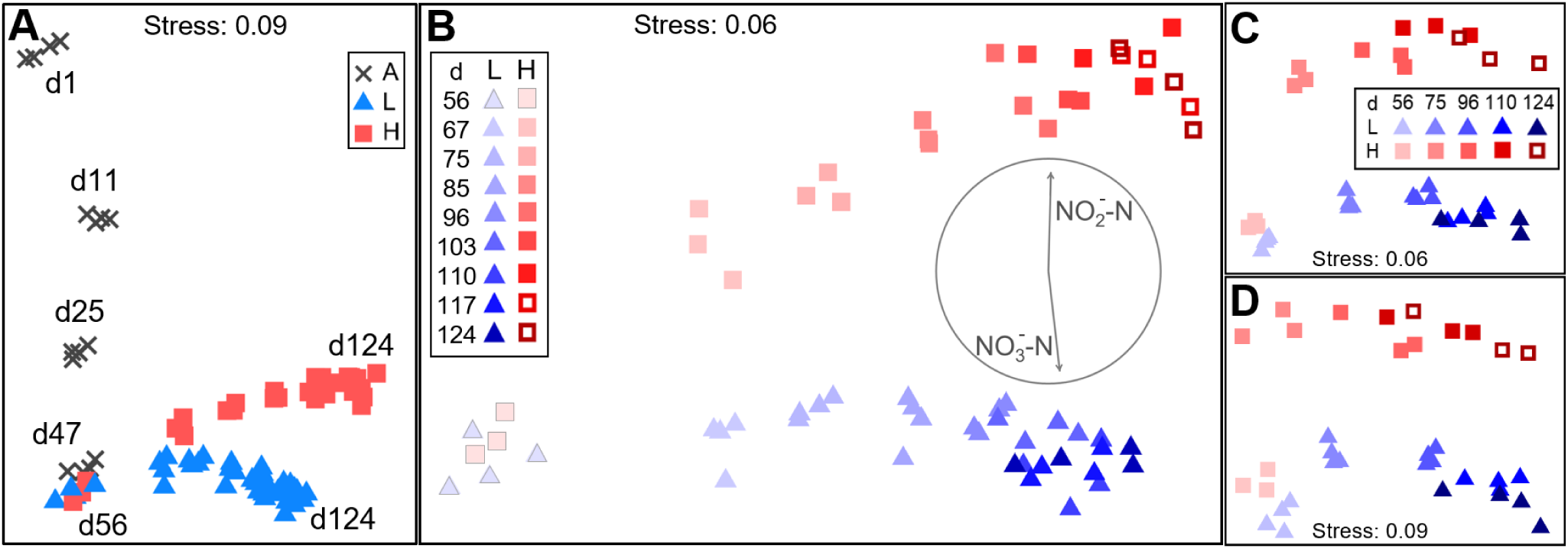
Community structure dynamics for bacterial taxa and functional genes evaluated through NMDS ordination (Bray-Curtis β-diversity). **(A**-**B)** 16S rRNA gene sequencing ASV level, (**C**) metagenomics sequencing genus level, and (**D**) metagenomics sequencing IP2G lowest gene level (same legend as **C**). Phases: A, acclimation (grey crosses, n = 4); L, low organic loading (blue triangles, n = 4); H, high organic loading (red squares, n = 3). Open red squares indicate the shift from high to low organic loading. Each point represents a different reactor for a given day. Days indicated with: (**A**) text within the panel for d1-d124; (**B**-**D**) decreasing brightness from d56-d124. Panel (**B**) includes Pearson’s correlation vectors of nitrite and nitrate in the effluent.

Patterns of α-diversity varied with time and differed among treatments (Fig. 1, Fig. S1). Press disturbed reactors displayed higher α-diversity (^2^D) for taxonomic datasets of both sequencing techniques employed (Fig. 1A-B). In contrast, undisturbed reactors displayed the highest α-diversity of functional genes at the end of the disturbance phase (Fig. 1C). Lower order compound α-diversity (^1^D) showed similar patterns, while richness (^0^D) did not show any differences among treatments (Fig. S1). After the shift from high to low organic loading, treatments continued to vary in terms of α-diversity, with ^2^D being significantly different for both the ASV (P_d124_ = 0.0046) and the IP2G gene datasets (P_d124_ = 0.0003) based on Welch’s ANOVA.

During the disturbance phase, temporal patterns of β-diversity showed bacterial communities clustering separately for low and high organic loading treatments regardless of the sequencing method employed for both taxa and genes (Fig. 2).

As per PERMANOVA, such differentiation was statistically significant from d56 onwards for all datasets, with non-significant PERMDISP results supporting the absence of heteroscedasticity (Table S2). Dissimilarity increased with time among replicates of both treatments across all datasets (Fig 2B-D). There was no significant effect of the shift from high to low organic loading with regards to β-diversity.

Similarly to observed diversity patterns, succession was evident after analysis of bacterial relative abundances. The 15 most abundant genera, assessed through 16S rRNA gene metabarcoding at the genus level, were different at each phase of the study (Fig. 3). Only six taxa were present in the top 15 genera throughout the entire study, and their relative abundances differed with experimental phase. Succession was also observed in metagenomics data, at both the bacterial genus taxonomic (Fig. S2) and the functional gene level (Fig. S3) (see details in supplementary information).

**Fig. 3.**
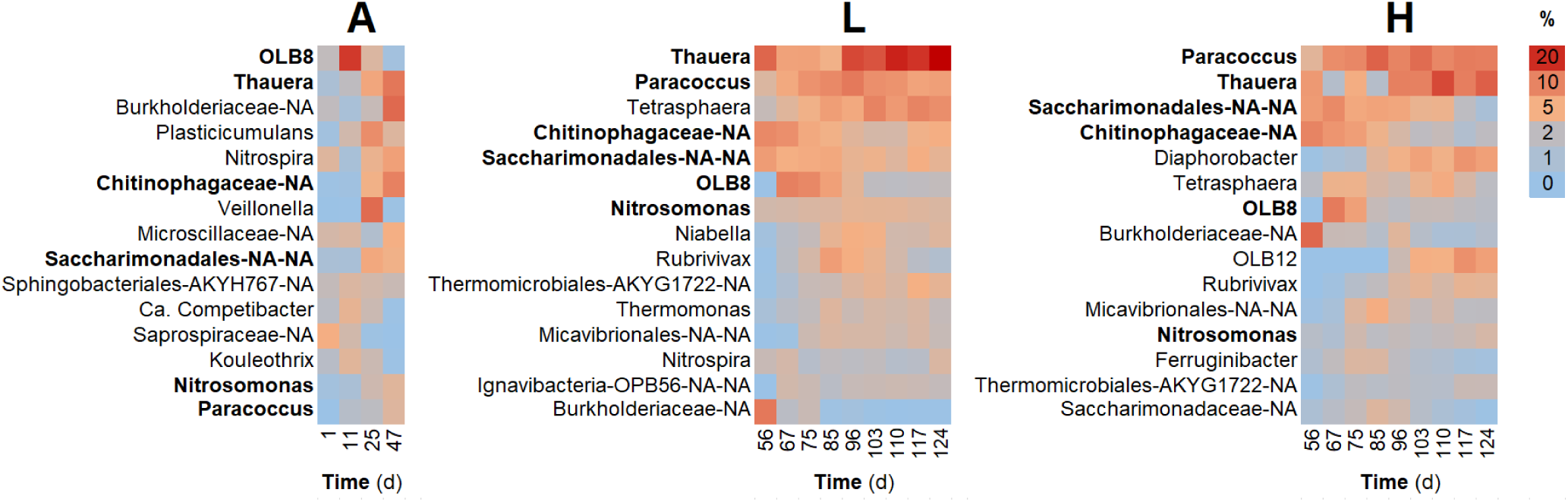
Heat maps of community structure dynamics for abundant bacterial taxa, assessed through 16S rRNA gene metabarcoding at the genus level. The 15 most abundant genera for each phase are shown. Genera that belong to the top 15 during all phases are shown in bold. Columns represent the average among reactors for a given phase and day. Phases: A, acclimation (left, n = 4); L, low organic loading (middle, n = 4); H, high organic loading (right, n = 3).

### Dynamics of ecosystem functions

Ecosystem function dynamics were described in detail elsewhere [44]. In short, there was a clear distinction between low and high organic loading reactors, in terms of nitrification and organic carbon removal, with the press disturbed reactors displaying partial nitrification with high NO_2_-N and COD concentrations in the effluent (Fig. S4). Following the shift from high to low organic loading, a transition towards recovery of the carbon and nitrite oxidation functions was observed, with a high variability across reactors (Fig. S4).

### Dynamics of bacterial community assembly mechanisms

Deterministic strength (DS), a metric that quantifies deterministic assembly, was generally higher for press disturbed reactors regardless of the sequencing method (Fig. 4) with a higher separation for taxa (Fig. 4A,D) than for genes (Fig. 4G). Overall, DS decreased with time, which was more marked for metagenomics-based taxa and genes (Fig. 4D-I). This trend coincided with a temporal increase in β-diversity dissimilarity, within and among treatments (Fig. 2). Deterministic strength values were mostly above 50% for ASV data (Fig. 4A). In contrast, most of the DS values for the metagenomics IP2G gene level dataset were below 50% (Fig. 4G). Further partition of the datasets into common (up to 99% accumulated reads) and rare (< 1% accumulated reads) fractions, showed that the common fraction (Fig. 4B,E,H) had a generally higher DS than the rare one (Fig. 4C,F,I). After the shift from high to low organic loading, similar DS values for both levels were observed only for the metagenomics gene data (Fig. 4G-I). Observed β-diversity was greater than expected for bacterial genera regardless of the sequencing method 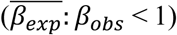, while an opposite pattern was seen for functional genes 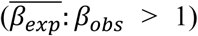 (Fig. 5). Detailed information of the parameters and outputs of the model are given in Tables S3-S5.

**Fig. 4.**
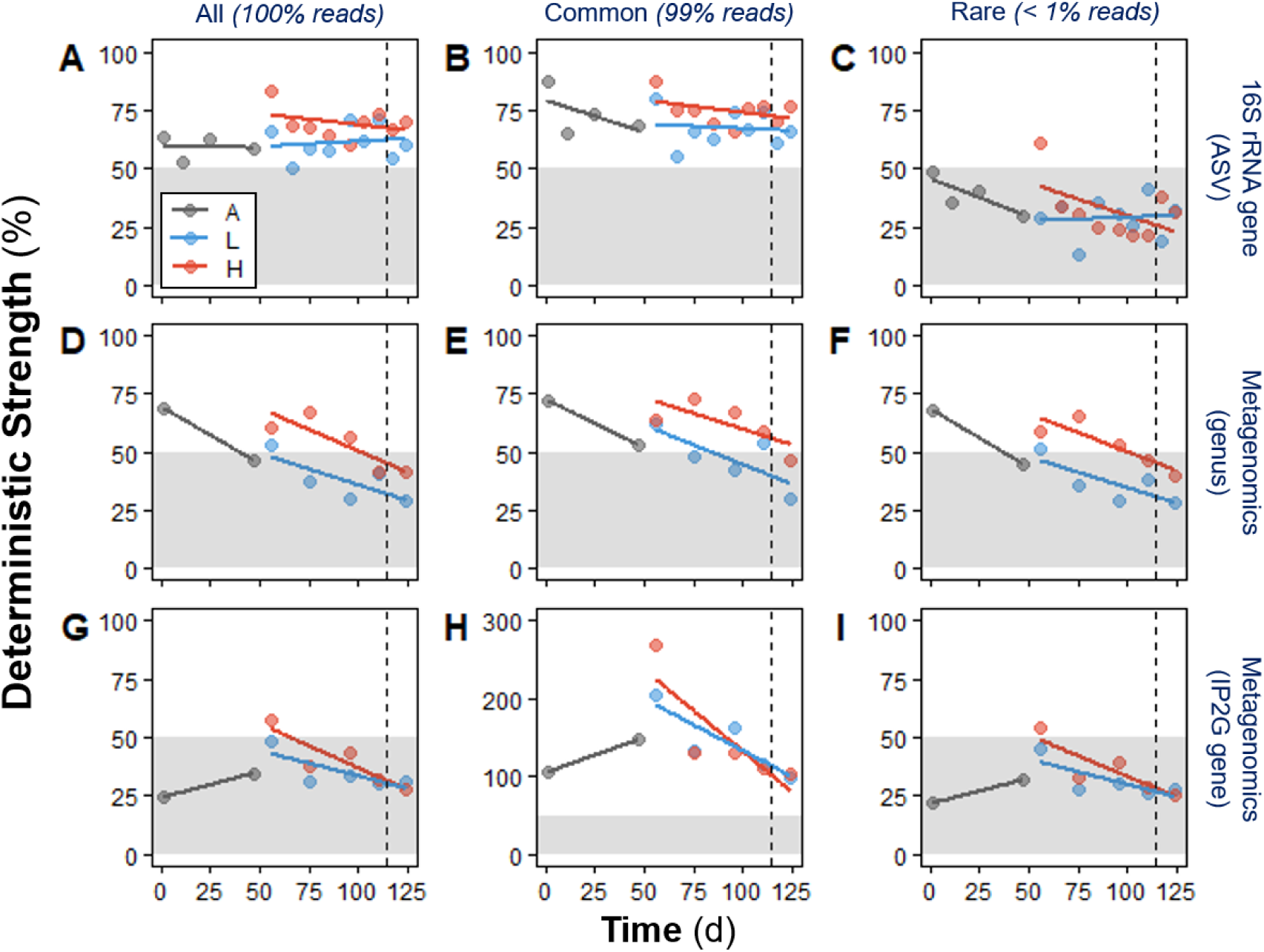
Deterministic strength (DS) temporal dynamics for bacterial taxa and functional genes, derived from null model analysis. (**A-C**) 16S rRNA gene sequencing ASV level. (**D-F**) Metagenomics sequencing genus level. (**G-I**) Metagenomics sequencing IP2G lowest gene level. Calculated using all taxa/genes (left panels: **A, D, G**), common taxa/genes (99% acc. reads; middle panels: **B, E, H**), and rare taxa/genes (<1% acc. reads; right panels: **C, F, I**). Phases: A, acclimation (grey); L, low organic loading (blue); H, high organic loading (red). Each point calculation involved all replicates (n = 4 for A and L, n =3 for H) of each phase evaluated. Lines represent linear regression fitting. Vertical dashed line indicates the shift from high to low organic loading. Zones of higher stochastic intensity within the panels are shaded in grey.

**Fig. 5.**
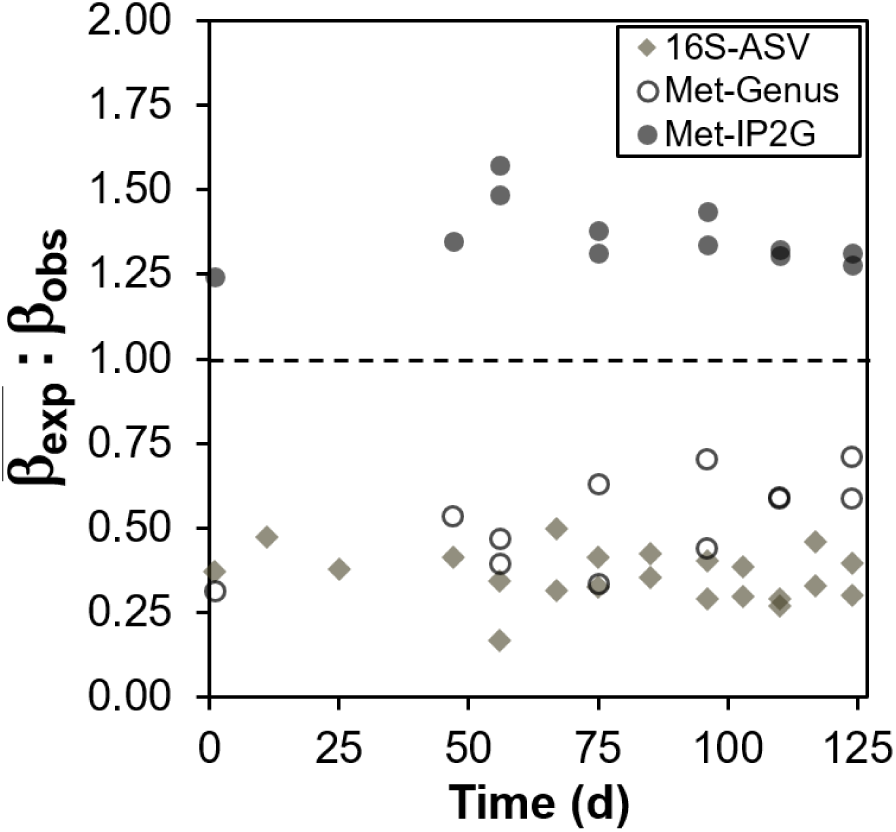
Expected to observed β-diversity ratio 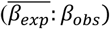, derived from null model analysis. Values below one (dashed line) indicate that the observed β-diversity is greater than expected by chance under the null model, while values above one indicate the opposite. Rhombuses represent 16S rRNA gene data at the bacterial ASV level. Circles represent shotgun metagenomics data at the bacterial genus (open circles) and lowest IP2G gene (closed circles) levels. Calculated using all taxa/genes. Each point calculation involved all replicates (n = 4 for A and L, n = 3 for H) of each phase evaluated.

## Discussion

### Function and community stabilization during acclimation phase

Sludge acclimation was a necessary transient stage in terms of ecosystem function and community structure. This period allowed for important functions like nitrification and organic carbon removal to stabilize across reactors (Fig. S4), as previously suggested [59]. We observed that α-diversity for both bacterial genera and functional genes decreased during acclimation (Fig. 1), which is coherent with the shift in β-diversity clusters (Fig. 2A). A decrease in bacterial diversity in bioreactors is expected, due to radically changing conditions when communities are moved from a full-scale plant to a laboratory setting, and has been observed for both anaerobic [60] and aerobic [21] bioreactors.

### Disturbance leads to different community function and structure

There was a clear distinction in function and community structure, in terms of α-and β-diversity, between low and high organic loading reactors. We previously reported on the effect of the high organic loading press disturbance on organic carbon removal and nitrite oxidation processes in this experiment [44]. With regards to community structure, the dominant genera of the acclimation phase gave way to two separate clusters with different dominant genera at the end of the two treatment levels (Fig. 3, Fig. S2). Bacterial successional dynamics have been described before for activated sludge systems, but studies often present analysis at the phylum or class levels of taxonomy [26, 27, 61]. Analysis of broad taxonomic categories may hide important assembly mechanisms operating at lower taxonomic levels [62], the reason why in this study we present successional changes at the genus level. We further evaluated succession of trait-complexes [63], identifying patterns that suggest differential gene investment across organisms at different stages of the study and organic loading levels. Traits like cell motility and cell wall were enriched in low organic loading reactors, while traits of replication and repair prevailed in high organic loading ones, suggesting the presence of community-level tradeoffs under disturbance similar to the ones described by life-history strategy theory [51]. Additionally, some of the press disturbed reactors recovered the carbon and nitrite oxidation function after returning to low organic loading conditions for 14 days (Fig. S1). However, their α− and β-diversity with respect to either taxonomy or functional genes did not recover (Figs. 1 and 2), which supports the insurance hypothesis [64] and functional redundancy [65] since an altered community was able to still provide the functions of the original one.

### Higher α-diversity does not translate into better function

The high organic loading disturbance led to inhibition of the nitrite oxidation function in these reactors, coincidently with a marked community differentiation from the low organic loading reactors in terms of β-diversity (Fig. 2B). However, we also found higher taxonomic α-diversity (^2^D_ASV_, ^2^D_Genus_) among the high organic loading reactors (Fig. 1), which means that the relative abundances were more evenly distributed among taxa for such communities. On the contrary, prior studies reported a decrease in Shannon diversity [66, 67] and Hill numbers [61] at higher organic loading. Since the low organic loading reactors displayed better COD removal and complete nitrification with almost no residual NH_3_-N or NO_2_^-^-N, it was expected that they would harbour more diverse communities, as more even communities were reported to have better functionality [68]. We did observe higher functional gene [42] α-diversity (^2^D_IP2G_, Fig. 1C) for the low organic loading reactors towards the end of the study, which is similar to the opposing trends of taxonomic and functional gene α-diversity previously reported after autoclave sterilization of soil microbial communities [69]. However, other studies using sludge bioreactors reported a negative relationship between bacterial richness and bioreactor performance [70], found no consistent relationships between α-diversity (richness, evenness) and the removal of endocrine disrupting compounds [71], and reported a negative correlation between ^2^D and COD removal but a positive one with NO_2_^-^-N generation [21]. This suggests that relationships between taxonomic α-diversity and function are not universal but more dependent on the system being evaluated and its particular conditions. Furthermore, it was proposed that α-diversity should be evaluated in context to draw meaningful insights about community mechanisms [72], while it was also suggested that changes in community assembly might be shaping the relationships between diversity and function [73]. Thus, for activated sludge systems different disturbances may elicit different relationships, at the taxonomic and functional gene levels, between α-diversity and important functions like nitrification.

### Community assembly processes differ for taxa and genes, as well as for common and rare fractions

In terms of relative deterministic strength, press disturbed reactors showed a stronger role of deterministic processes for all three types of datasets evaluated (Fig. 4), likely due to the selective pressure via environmental filtering [74] imposed by the disturbance. Deterministic mechanisms dominated bacterial community assembly at the ASV taxonomic level across all reactors (DS > 50%) and were stronger under high organic loading (Fig. 4A). This agrees with previous studies of press disturbance in mesocosm sludge bioreactors using a micropollutant (3-chloroaniline), which reported higher similarity among disturbed reactors compared to the similarity among control reactors via 16S rRNA gene amplicon analysis [75, 76], although without quantifying assembly mechanisms. Partitioning the contribution of common (99% reads accumulated) and rare (<1% reads accumulated) portions of the data revealed that the common fraction was driving the overall deterministic assembly mechanisms (Fig. 4B), while the rare portion displayed high stochasticity (Fig. 4C). These findings are in agreement with prior studies that reported higher variability for rare taxa compared to common taxa in full-scale and mesocosm bioreactor sludge systems [26-29], albeit without the use of null model analysis. Stochastic processes of ecological drift can operate in the portion of taxa at low relative abundances [20], while the overall community assembly is mainly deterministic. Further, a recent study on granular biofilm reactors using one simple carbon source also reported stronger selection for abundant taxa and higher stochastic assembly via drift for low-abundant taxa [38], by using a different null model analysis that takes into account phylogeny but does not take advantage of replicated designs [77]. All these prior studies employed 16S rRNA gene metabarcoding with OTU type of clustering for their community analyses. Ours is the first study to infer community assembly mechanisms from ASV data, which has several benefits over traditional OTU clustering [48]. Although classification criteria for rare/common fractions are arbitrary [30, 78], there is consistency between our community assembly observations from 16S rRNA gene metabarcoding data and the current literature. Nonetheless, further research is needed to evaluate whether the rare fraction is biologically active [79].

Metagenomics data also revealed higher deterministic strength at high organic loading for both taxa (Fig. 4D) and genes (Fig. 4G), albeit with lower DS values compared to those for the ASV dataset. This concurs with the higher deterministic assembly reported for sludge microcosm reactors that were press disturbed with 3-chloroaniline and compared to undisturbed ones, via null model analysis on metagenomics genus-level data [21]. Further, most of the DS points for the overall IP2G gene dataset fell below 50% (Fig. 4G), implying that stochastic mechanisms of assembly were stronger for genes in comparison with taxa. Separate evaluation of the common and rare fractions in terms of DS showed that deterministic assembly mechanisms were stronger in the common fraction (Fig. 4E,H) while the rare portion was more influenced by stochasticity (Fig. 4F,I), which is similar to what we observed for ASV data. Conversely, analysis of metagenomics data revealed the overall community assembly to be driven by the rare fraction, particularly for the functional gene dataset (Fig. 4I). Further, beta diversity was always greater than expected for bacterial taxa under the null model regardless of the sequencing method (Fig. 5, Tables S3-S5), suggesting that taxa tended to be more aggregated within replicate bioreactors than expected by chance. Aggregation can be explained by processes of habitat filtering [80] due to the recurrent conditions at each low and high organic loading level, as well as by dispersal limitation [81], which is a condition of the experimental design using reactors as closed systems without immigration. However, the opposite was shown for the functional gene dataset where the mean expected beta diversity of individual IP2G genes under the null model was higher than the observed beta diversity, i.e., replicate bioreactors were more similar than expected by chance (Fig. 6). These results highlight that functional gene data can indicate patterns of dominant stochastic versus deterministic mechanisms of community assembly that differ from patterns obtained after analysis of taxon data.

### Sequencing methods, diversity and assembly mechanisms: what is consistent and what is not

Using two different community profiling approaches on the same samples we found that press disturbance favoured deterministic assembly mechanisms, where bioreactor bacterial communities at low and high organic loading levels clearly separated into two different clusters in terms of β-diversity for taxa and genes. Both methods identified a greater influence of deterministic assembly mechanisms in the common fraction of the community, whereas stochasticity was more important in the low abundance fraction. These complementary analyses suggest one should be able to assess the effect of a press disturbance on both β-diversity and the overall assembly mechanisms of microbial communities, regardless of the sequencing approach employed and whether the focus on taxa or functional genes.

However, methodologies were inconsistent when probing the relative contributions of deterministic and stochastic mechanisms of community assembly, with metagenomics data displaying higher stochasticity for genera compared to when ASVs from 16S rRNA gene metabarcoding were used. This was not an effect of assessing different levels of taxonomic resolution, as higher resolution levels were shown to be more conserved than lower ones [62], and therefore displayed assembly mechanisms that were more deterministic in nature. Using metagenomics data, functional genes were found to be more stochastically driven than taxa at the genus level. Further, there were differences in terms of which fraction of the community, common or rare, affected community assembly. The driver for 16S rRNA amplicon data was the common fraction and for metagenomics data, it was the rare fraction. Similarly, the effect of disturbance on α-diversity was different for genes and taxonomic data. The challenge of reconciling results from different sequencing methods has been recognized and requires further research [41]. Differences in assembly mechanisms can arise due to the distinct biological signature being evaluated when targeting a fraction of a specific DNA marker like the 16S rRNA gene, compared to using DNA of the whole meta-community as the basis of classification of taxa and functional genes. Currently, the 16S rRNA amplicon approach is expected to result in less ambiguity because fewer singletons are generated than with shotgun metagenomics. Not surprisingly, the majority of existing studies on microbial community assembly focused on taxonomic OTU datasets from 16S rRNA gene metabarcoding [17, 18, 33-38], while a few others evaluated functional data via GeoChip [22, 43]. To our knowledge, only one previous study used metagenomics data to interrogate assembly mechanisms in microbial communities [21], despite the fact that the trend in microbial community analysis is moving towards metagenomics with the advent of faster, PCR-independent, long-read sequencing [82].

### Concluding remarks

The joint evaluation of assembly mechanisms, community structure and function of bacterial taxa and functional genes provided in-depth understanding of the response of complex microbial systems to a doubling of the organic loading rate. This press disturbance altered community function, structure and assembly mechanisms. Disturbance had an effect not only on community function but also on its functional potential, emphasising the relevance of assessing communities of organisms together with communities of genes. We highlighted that higher α-diversity does not always imply better function and that press disturbance favoured deterministic assembly mechanisms. Community assembly was found to have a stronger deterministic component among the common fraction, whereas the role of stochastic mechanisms was higher for the less abundant portion of the community. Also, reactors who recovered functions after returning to low organic loading conditions maintained different α- and β-diversities compared to reactors that had not been press-disturbed, in terms of both taxonomic and functional genes, showing that resilience based on community function does not necessarily translate into resilience based on community structure. Finally, we urge caution when assessing microbial community assembly mechanisms, as results can vary depending on the approach (16S rRNA gene metabarcoding, shotgun metagenomics taxonomic or functional gene community profiling) and whether the focus is on taxa or genes. Not only can genes suggest different dominance patterns of stochastic versus deterministic mechanisms of community assembly compared to taxa, but the fraction of the community driving such assembly mechanisms – common versus rare – can also differ.

## Acknowledgements

This research was supported by the Singapore National Research Foundation and Ministry of Education under the Research Centre of Excellence Program. We thank DI Drautz-Moses for her support with the 16S rRNA gene amplicon and metagenomics library preparation and sequencing pipelines employed. WX Phua is acknowledged for her support with bioreactors operation. We thank TJ Qiang and NABA Latiff for their assistance with molecular work. ES was partially supported by a Fulbright Fellowship.

## Author Contributions

ES and SW conceived the study. ES designed the experiment. SW obtained the funding for the study. ES performed the experiments and the 16S rRNA gene bioinformatics analyses. FC contributed the metagenomics bioinformatics analyses. ES interpreted the data and elaborated the main arguments in the manuscript. ES and SW wrote the manuscript.

## Data availability

DNA sequencing data are available at NCBI BioProjects PRJNA559245. See supplementary information for details about community structure dynamics based on bacterial genera and functional genes, as well as arguments discouraging the use of richness as an indicator of α-diversity.

