## Supplementary Information for "Press disturbance unveils different community structure, function and assembly of bacterial taxa and functional genes in mesocosm-scale bioreactors"

#### Supplementary text

*Bacterial genus-level structure dynamics*

Analysis of changes in the 100 most abundant genera also revealed bacterial succession in different clusters (Fig. S2). Taxa that prevailed in the sludge inoculum (d1) like *Ca.* Accumulibacter, *Desulfovibrio,* and *Mycobacterium* had their relative abundances reduced to very low levels during the remainder of the study. Different genera dominated in samples at the end of the acclimation phase (d47) and at the beginning of the experimental phase (d56), for example, *Bosea*, *Acidovorax*, *Nakamurella*, *Comamonas*, and *Microlunatus*. The organisms prevailing in disturbed and undisturbed reactors also varied from d75 onwards. *Tetrasphaera*, *Nitrosomonas*, and *Thauera* preferred the low organic loading conditions, whereas genera like *Gemmatimonas*, *Propionibacterium*, and *Kineosphaera* prevailed at high organic loading. After the switch back from high to low organic loading, *Ca.* Competibacter and *Caldilinea* showed an increase in abundance to levels comparable to the ones found in the inoculum.

*Bacterial functional gene dynamics*

Functional potential analysis displayed a succession of functional genes in differentiated groups (Fig. S3), similar to the succession we observed for taxa (Fig. 3, Fig. S2). We focused on trait complexes which are a product of the expression of multiple true traits [1], grouping different sets of genes into categories. Within the COG database (Fig. S3A) trait categories like lipid transport and metabolism, energy production and conversion, and secondary metabolites biosynthesis, transport and catabolism were the ones prevailing at the sludge inoculum. After the acclimation phase abundant genes related to transcription, chromatin structure and dynamics, and intracellular trafficking. Interestingly, trait complexes related to transport and metabolism of carbohydrates, amino acids and nucleotides increased for both high and low organic loading treatments at an early stage (d75), but then gave way to other functional genes. Genes encoding for cell motility and the cell envelope prevailed in low organic loading reactors afterwards, whereas replication, recombination and repair, and carbohydrate transport metabolism did so in high organic loading reactors. Inorganic ion transport and metabolism genes were enriched in both treatments from d110 onwards. Functional gene data from other gene databases exhibited similar successional trends (Figs. S3B-D) and also allowed us to identify some trait complexes that were enriched after the switch from high to low organic loadings, like genes related to cell wall and capsule (Fig. S3C) and receptor activity (Fig. S3D).

*Richness is not recommended as $\alpha$-diversity metric for microbial community studies*

With regards to richness ($^{0}D$) we acknowledge that this is not a robust indicator of microbial community diversity due to current sequencing and bioinformatics limitations [2]. For example, the increase in $^{0}D_{Genus}$ diversity for low organic loading reactors during the disturbance phase (Fig. S1D) likely means that more genera increased in abundance above the limit of detection, as new organisms could not be incorporated since our system was closed to immigration. The latter implies that these taxa were already in the reactors but at levels below the detection limit. This phenomenon could happen during any study on complex microbial communities leading to spurious patterns without ecological meaning. It is, therefore, not recommended to draw conclusions based on bacterial richness dynamics.


### Supplementary Figures

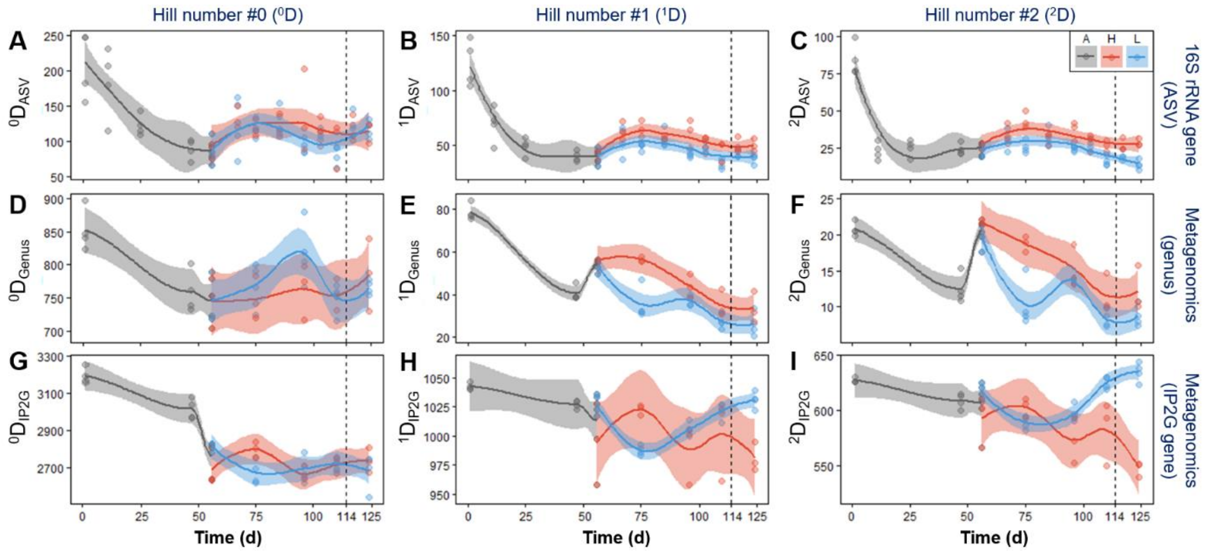


**Fig. S1**. Temporal dynamics of true $\alpha$-diversity Hill numbers ($^0$D, $^1$D, $^2$D) for bacterial taxa and functional genes. (**A**-**C**) 16S rRNA gene sequencing ASV level, (**D**-**F**) metagenomics sequencing genus level, and (**G**-**I**) metagenomics sequencing IP2G lowest gene level. Each point represents a different reactor for a given day. Phases: A, acclimation (grey, n = 4); L, low organic loading (blue, n = 4); H, high organic loading (red, n = 3). Vertical dashed line indicates the shift from high to low organic loading. Lines refer to polynomial regression fitting, while shaded areas represent 95% confidence intervals.

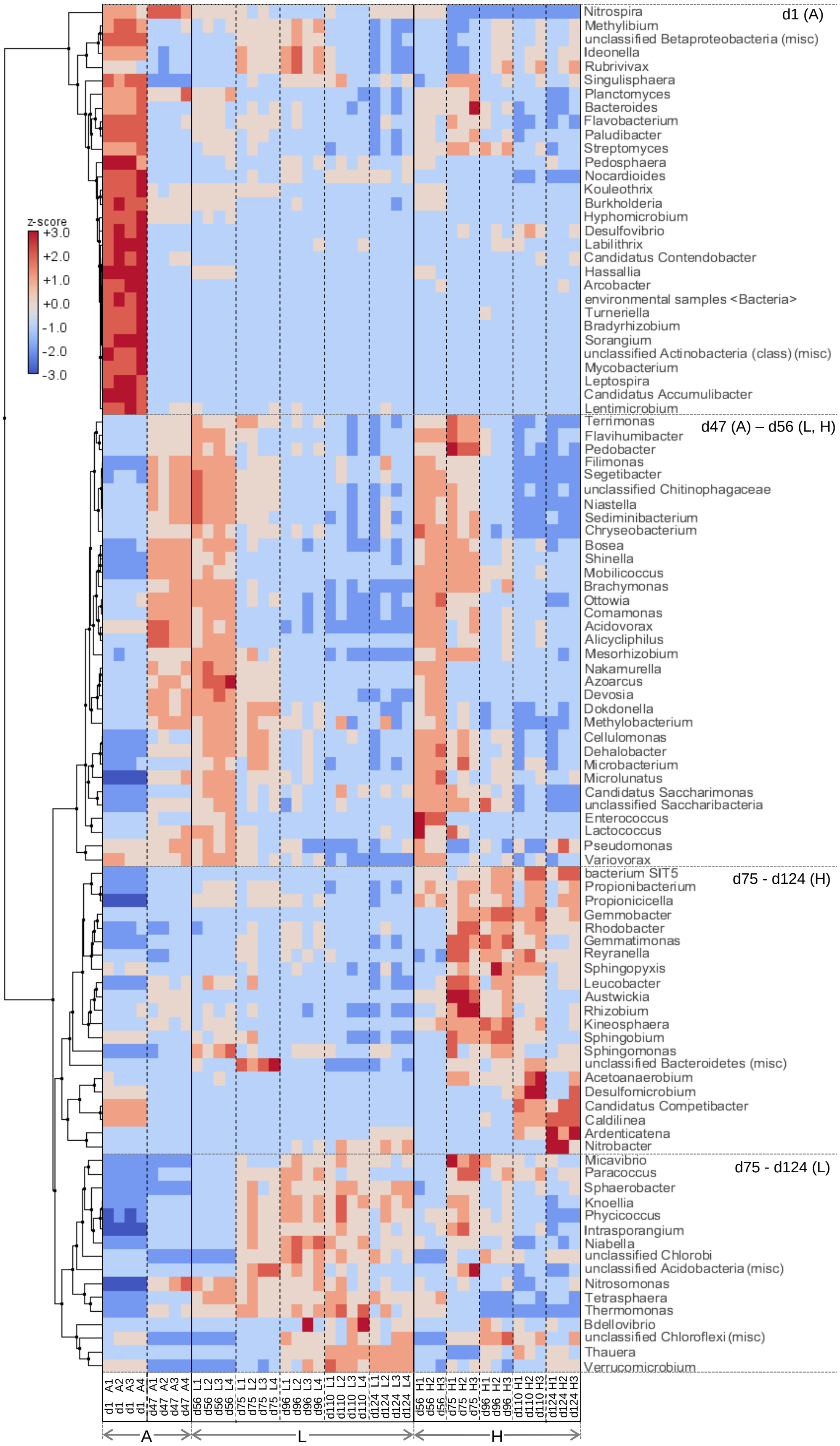


**Fig. S2.** Temporal comparisons of most abundant genera help discern successional clusters. Clustered heat map for the 100 most abundant genera in reactors over time. Z-scores denote how many standard deviations of the mean (assigned reads per genera, across all samples) each sample contains. Column legend represents day number and replicate reactors. Phases: A, acclimation (n = 4); L, low organic loading (n = 4); H, high organic loading (n = 3). Rectangles highlight groups of taxa prevailing at different phases. Dashed vertical lines separate time points within the same phase. Dashed horizontal lines separate the four largest clusters.

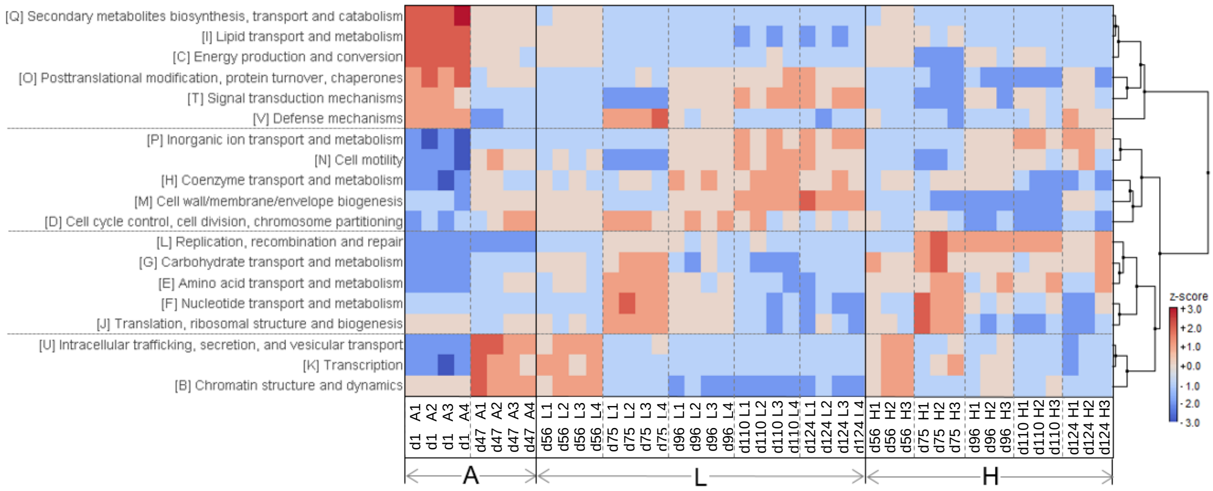


**Fig. S3A.** Successional clusters of functional gene potential. Clustered heat map for the 19 most abundant COG trait complex categories (>10000 reads) in reactors over time. Z-scores denote how many standard deviations of the mean (assigned reads per gene category, across all samples) each sample contains. Column legend represents day number and replicate reactors. Phases: A, acclimation (n = 4); L, low organic loading (n = 4); H, high organic loading (n = 3). Rectangles highlight taxa groups prevailing at different phases. Dashed vertical lines separate time points within the same phase. Dashed horizontal lines separate the four biggest clusters.

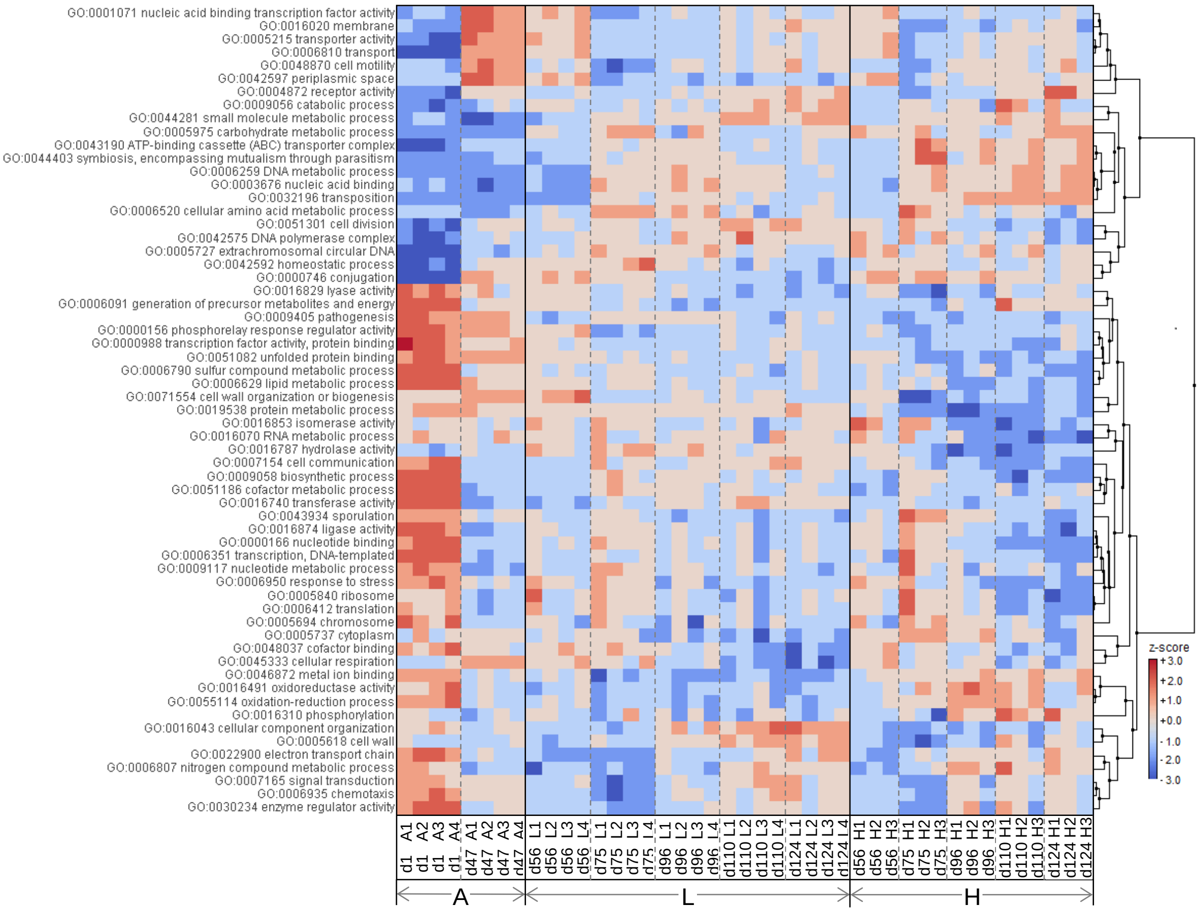


**Fig. S3B.** Clustered heat map for the 61 most abundant IP2G trait complex categories (>10000 reads) in reactors over time. Z-scores denote how many standard deviations of the mean (assigned reads per gene category, across all samples) each sample contains. Column legend represents day number and replicate reactors. Phases: A, acclimation (n = 4); L, low organic loading (n = 4); H, high organic loading (n = 3). Rectangles highlight taxa groups prevailing at different phases. Dashed vertical lines separate time points within the same phase.

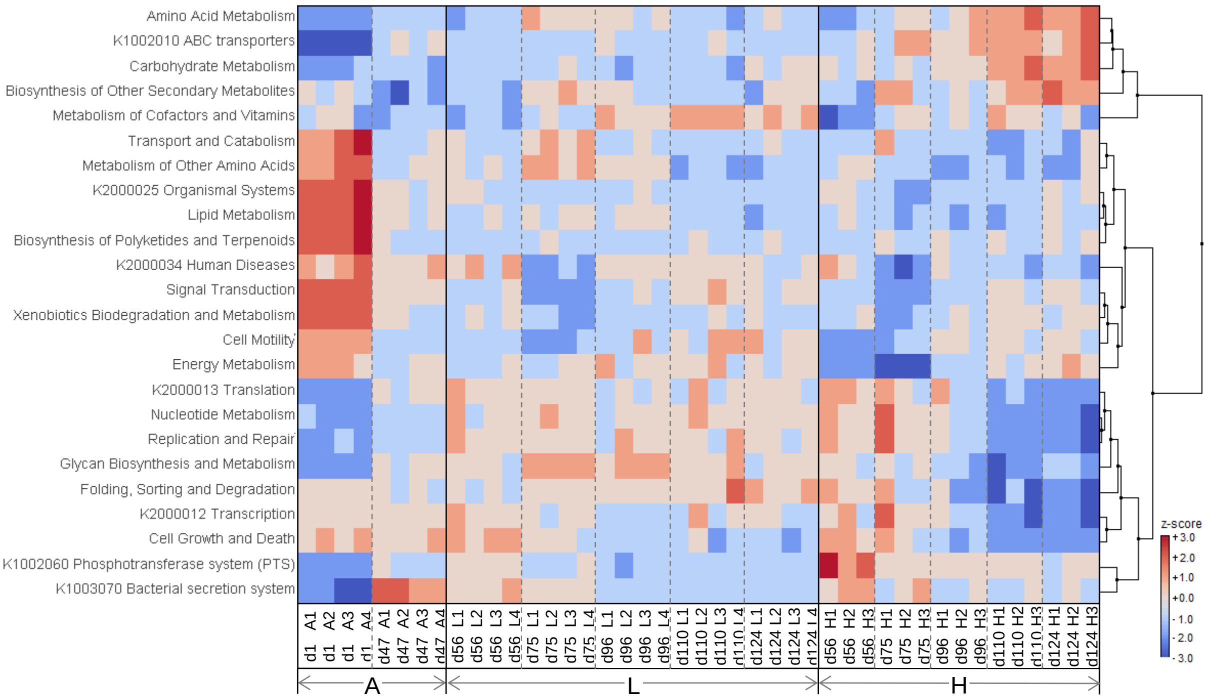


**Fig. S3C.** Clustered heat map for the 20 most abundant KEGG trait complex categories (>10000 reads) in reactors over time. Z-scores denote how many standard deviations of the mean (assigned reads per gene category, across all samples) each sample contains. Column legend represents day number and replicate reactors. Phases: A, acclimation (n = 4); L, low organic loading (n = 4); H, high organic loading (n = 3). Rectangles highlight taxa groups prevailing at different phases. Dashed vertical lines separate time points within the same phase.

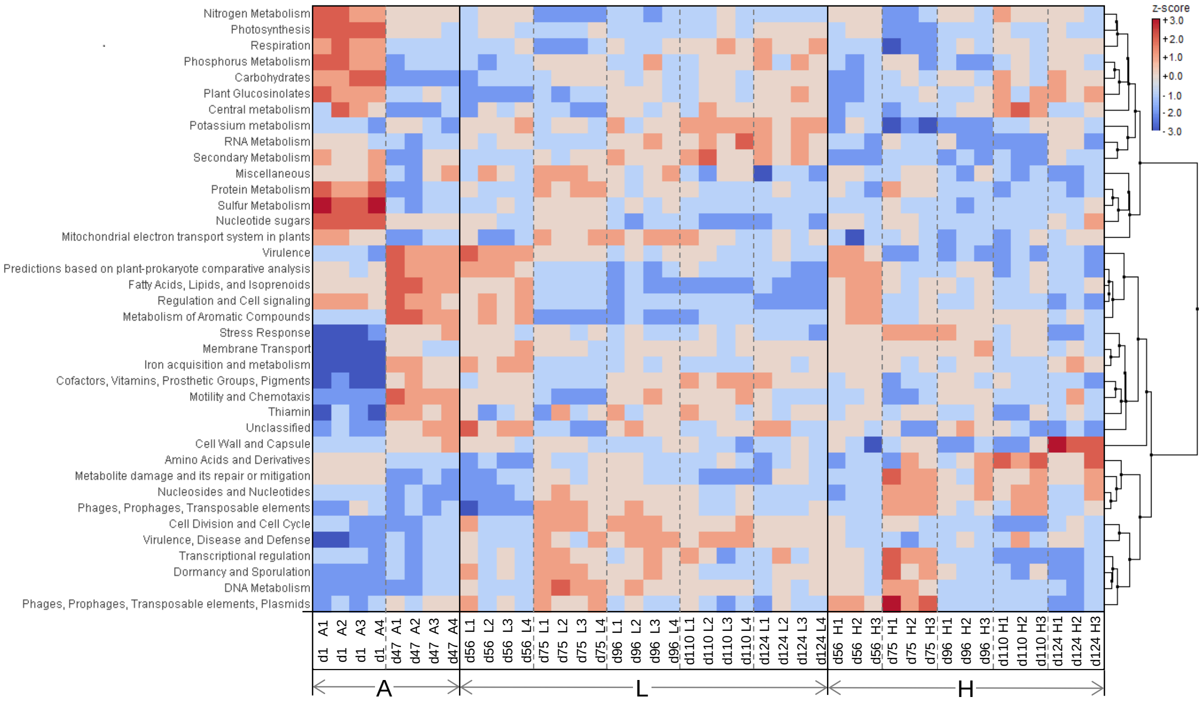


**Fig. S3D.** Clustered heat map for the 38 most abundant SEED trait complex categories (>10000 reads) in reactors over time. Z-scores denote how many standard deviations of the mean (assigned reads per gene category, across all samples) each sample contains. Column legend represents day number and replicate reactors. Phases: A, acclimation (n = 4); L, low organic loading (n = 4); H, high organic loading (n = 3). Rectangles highlight taxa groups prevailing at different phases. Dashed vertical lines separate time points within the same phase.

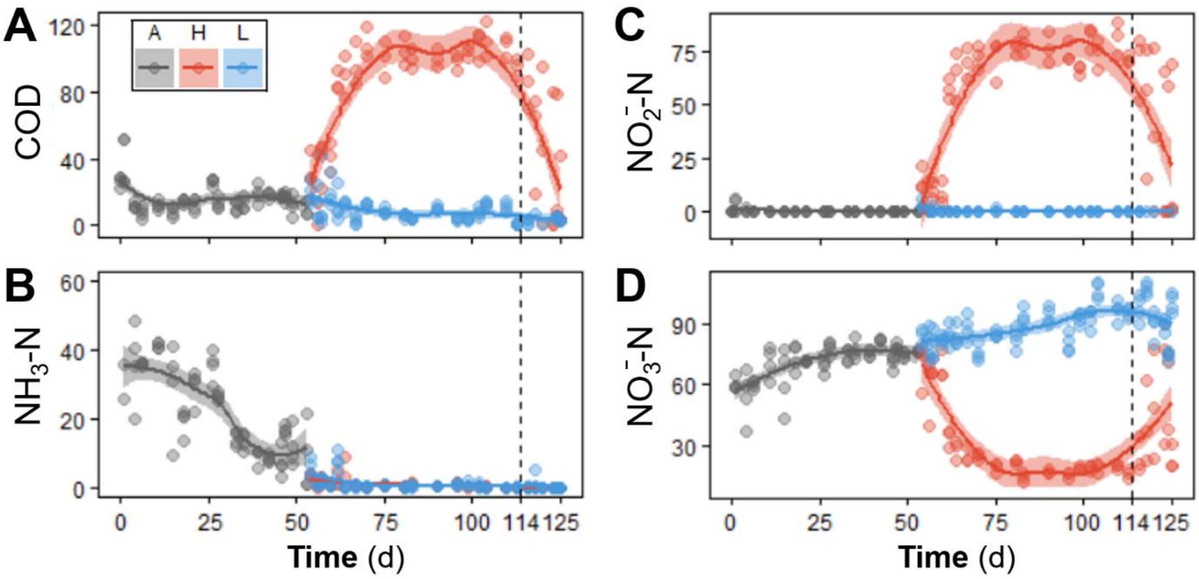


**Fig. S4.** Temporal effluent concentrations (mg $L^{-1}$) at the end of a reactor cycle, adapted from [3]. (**A**) Organic carbon as soluble chemical oxygen demand (COD), and (**B**) ammonia, (**C**) nitrite, and (**D**) nitrate as nitrogen. Each point represents a different reactor for a given day. Phases: A, acclimation (grey, n = 4); L, low organic loading (blue, n = 4); H, high organic loading (red, n = 3). Vertical dashed line indicates the shift from high to low organic loading. Lines display polynomial regression fitting, while shaded areas represent 95% confidence intervals.

### Supplementary Tables

**Table S1.** Reactor parameters and influent synthetic wastewater characteristics per phase [3].

| Day | Phase† | *n* | F:M* [mg-COD mg-TSS$^{-1}$ d$^{-1}$] | C:N [mg-COD mg-TKN$^{-1}$] | COD [mg L$^{-1}$] | TKN [mg L$^{-1}$] | TSS [mg L$^{-1}$] | SRT [d] |
|---|---|---|---|---|---|---|---|---|
| 1-53 | Acclimation | 4 | 0.21 (0.08) | 3.5 (0.7) | 374 (106) | 105 (27) | 1934 (502) | 18.7 (0.7) |
| 54-127 | Low organic loading | 4 | 0.19 (0.05) | 3.5 (0.3) | 323 (24) | 92 (3.6) | 1727 (251) | 12.8 (0.4) |
| 54-113 | High organic loading‡ | 3 | 0.36 (0.11) | 6.3 (0.9) | 629 (67) | 100 (19) | 1943 (476) | 8.2 (1.4) |
| 114-127 | High-to-low organic loading‡ | 3 | 0.19 (0.06) | 3.6 (0.3) | 326 (19) | 90 (2.2) | 1774 (256) | 12.4 (1.0) |

* Average values, including standard deviation of the mean (s.d.m.) between parentheses.
† Each phase represents *n* replicates of independent 5-L reactors. Samples were generated 2-3 times per week.
‡ These two phases involved the same reactors, where organic loading was changed from high to low on d114.
F:M, food-to-biomass ratio
C:N, carbon-to-nitrogen ratio
COD, chemical oxygen demand
TKN, total Kjeldahl nitrogen
TSS, total suspended solids
SRT, solids residence time

**Table S2.** Multivariate tests for relative abundances of bacterial communities (taxa and functional genes). Disturbance and time set as factors. BC dissimilarity metric used. P-values adjusted at a FDR of 5%.

| Sequencing method | Factor* | No of levels | *n*‡ | df§ res | PERMANOVA† | | PERMDISP† | |
|---|---|---|---|---|---|---|---|---|
| | | | | | F | P (MC)¶ | F | P (perm) |
| 16S rRNA gene metabarcoding<br>ASV taxonomic level | t (d1-d47) | 2 | 4 | 6 | 30.511 | **0.0036**‖ | 180.44 | 0.0576 |
| | OL (d56) | 2 | 4, 3 | 5 | 1.6269 | 0.2942 | 0.2582 | 0.9006 |
| | OL (d75) | 2 | 4, 3 | 5 | 9.9127 | **0.0086** | 0.4442 | 0.5912 |
| | OL (d96) | 2 | 4, 3 | 5 | 8.9456 | **0.0098** | 2.7755 | 0.4470 |
| | OL (d110) | 2 | 4, 3 | 5 | 10.429 | **0.0080** | 1.5584 | 0.4715 |
| | OL (d124) | 2 | 4, 3 | 5 | 12.032 | **0.0078** | 0.5991 | 0.6363 |
| Metagenomics shotgun sequencing<br>Genus taxonomic level | t (d1-d47) | 2 | 4 | 6 | 193.11 | **0.0036** | 8.34 | 0.1948 |
| | OL (d56) | 2 | 4, 3 | 5 | 3.56 | 0.0576 | 0.0597 | 0.9375 |
| | OL (d75) | 2 | 4, 3 | 5 | 15.2 | **0.0080** | 1.5199 | 0.4715 |
| | OL (d96) | 2 | 4, 3 | 5 | 13.1 | **0.0078** | 0.8222 | 0.6532 |
| | OL (d110) | 2 | 4, 3 | 5 | 14.8 | **0.0054** | 0.2168 | 0.8984 |
| | OL (d124) | 2 | 4, 3 | 5 | 11.5 | **0.0080** | 0.7351 | 0.7343 |
| Metagenomics shotgun sequencing<br>IP2G lowest gene level | t (d1-d47) | 2 | 4 | 6 | 24.651 | **0.0048** | 29.441 | 0.0576 |
| | OL (d56) | 2 | 4, 3 | 5 | 1.8437 | 0.2292 | 0.0831 | 0.9006 |
| | OL (d75) | 2 | 4, 3 | 5 | 6.2232 | **0.0180** | 6.943 | 0.1097 |
| | OL (d96) | 2 | 4, 3 | 5 | 6.9173 | **0.0174** | 2.3795 | 0.4715 |
| | OL (d110) | 2 | 4, 3 | 5 | 7.14 | **0.0168** | 0.0006 | 0.9375 |
| | OL (d124) | 2 | 4, 3 | 5 | 6.3847 | **0.0211** | 0.8520 | 0.5912 |

* Factors (levels): t, time (d1 - d47) and OL, organic loading (low - high).
† Number of permutations used was 9,999
‡ Number of replicates per level
§ Degrees of freedom of the residual
¶ Approximate P-value from Monte Carlo permutations
‖ In bold, significant P-values after correction for multiple comparisons at a False Discovery Rate of 5%, using Benjamini-Hochberg's method.

**Table S3.** Parameter output from null model analysis on the ASV-level dataset (16S rRNA gene metabarcoding).

| **P**[†] | ***n***[‡] | **d**[§] | **All ASVs** (100% reads)[¶] | | | | | | **Common ASVs** (up to 99% acc. reads) | | | | | | **Rare ASVs** (less than 1% acc. reads) | | | | | |
|---|---|---|---|---|---|---|---|---|---|---|---|---|---|---|---|---|---|---|---|---|
| | | | $\gamma_{obs}$ | $\overline{\alpha_{obs}}$ | $\beta_{obs}$ | $\overline{\beta_{exp}}$ | **DS** (%) | $\overline{\beta_{exp}}:\beta_{obs}$ | $\gamma_{obs}$ | $\overline{\alpha_{obs}}$ | $\beta_{obs}$ | $\overline{\beta_{exp}}$ | **DS** (%) | $\overline{\beta_{exp}}:\beta_{obs}$ | $\gamma_{obs}$ | $\overline{\alpha_{obs}}$ | $\beta_{obs}$ | $\overline{\beta_{exp}}$ | **DS** (%) | $\overline{\beta_{exp}}:\beta_{obs}$ |
| A | 4 | 1 | 436 | 208 | 0.524 | 0.194 | 63.0 | 0.37 | 224 | 143 | 0.364 | 0.047 | 87.2 | 0.13 | 212 | 65 | 0.692 | 0.356 | 48.5 | 0.51 |
| | | 11 | 348 | 183 | 0.476 | 0.226 | 52.6 | 0.47 | 235 | 146 | 0.379 | 0.133 | 64.8 | 0.35 | 113 | 37 | 0.677 | 0.435 | 35.7 | 0.64 |
| | | 25 | 214 | 122 | 0.430 | 0.163 | 62.2 | 0.38 | 170 | 109 | 0.357 | 0.097 | 72.9 | 0.27 | 44 | 13 | 0.710 | 0.423 | 40.5 | 0.60 |
| | | 47 | 148 | 89 | 0.400 | 0.166 | 58.6 | 0.41 | 127 | 83 | 0.344 | 0.109 | 68.3 | 0.32 | 21 | 6 | 0.738 | 0.517 | 29.9 | 0.70 |
| L | 4 | 56 | 150 | 85 | 0.432 | 0.148 | 65.7 | 0.34 | 125 | 79 | 0.370 | 0.074 | 79.9 | 0.20 | 25 | 7 | 0.740 | 0.526 | 28.9 | 0.71 |
| | | 67 | 216 | 123 | 0.431 | 0.214 | 50.3 | 0.50 | 184 | 114 | 0.383 | 0.171 | 55.5 | 0.45 | 32 | 10 | 0.703 | 0.465 | 33.9 | 0.66 |
| | | 75 | 191 | 118 | 0.382 | 0.159 | 58.5 | 0.42 | 177 | 114 | 0.355 | 0.122 | 65.6 | 0.34 | 14 | 4 | 0.732 | 0.638 | 12.9 | 0.87 |
| | | 85 | 232 | 129 | 0.445 | 0.188 | 57.7 | 0.42 | 204 | 121 | 0.407 | 0.152 | 62.7 | 0.37 | 28 | 8 | 0.723 | 0.468 | 35.3 | 0.65 |
| | | 96 | 158 | 92 | 0.419 | 0.121 | 71.1 | 0.29 | 151 | 90 | 0.406 | 0.105 | 74.2 | 0.26 | 7 | 2 | 0.714 | 0.499 | 30.2 | 0.70 |
| | | 103 | 197 | 111 | 0.438 | 0.168 | 61.6 | 0.38 | 185 | 108 | 0.418 | 0.141 | 66.3 | 0.34 | 12 | 4 | 0.667 | 0.498 | 25.3 | 0.75 |
| | | 110 | 159 | 80 | 0.497 | 0.144 | 71.1 | 0.29 | 149 | 78 | 0.480 | 0.125 | 73.9 | 0.26 | 10 | 3 | 0.750 | 0.441 | 41.2 | 0.59 |
| | | 117 | 213 | 124 | 0.419 | 0.192 | 54.2 | 0.46 | 193 | 118 | 0.387 | 0.150 | 61.2 | 0.39 | 20 | 6 | 0.725 | 0.586 | 19.2 | 0.81 |
| | | 124 | 200 | 117 | 0.414 | 0.164 | 60.4 | 0.40 | 181 | 112 | 0.383 | 0.130 | 65.9 | 0.34 | 19 | 6 | 0.711 | 0.485 | 31.7 | 0.68 |
| H | 3 | 56 | 132 | 88 | 0.333 | 0.056 | 83.1 | 0.17 | 120 | 83 | 0.308 | 0.040 | 87.0 | 0.13 | 12 | 5 | 0.583 | 0.226 | 61.3 | 0.39 |
| | | 67 | 197 | 132 | 0.330 | 0.104 | 68.4 | 0.32 | 178 | 125 | 0.298 | 0.074 | 75.2 | 0.25 | 19 | 7 | 0.632 | 0.421 | 33.4 | 0.67 |
| | | 75 | 169 | 118 | 0.302 | 0.098 | 67.6 | 0.32 | 155 | 113 | 0.273 | 0.069 | 74.7 | 0.25 | 14 | 5 | 0.619 | 0.430 | 30.5 | 0.69 |
| | | 85 | 178 | 121 | 0.320 | 0.114 | 64.5 | 0.36 | 168 | 117 | 0.302 | 0.092 | 69.4 | 0.31 | 10 | 4 | 0.633 | 0.476 | 24.9 | 0.75 |
| | | 96 | 227 | 136 | 0.399 | 0.160 | 59.9 | 0.40 | 203 | 128 | 0.369 | 0.125 | 66.2 | 0.34 | 24 | 8 | 0.653 | 0.495 | 24.1 | 0.76 |
| | | 103 | 178 | 114 | 0.361 | 0.108 | 70.2 | 0.30 | 167 | 110 | 0.343 | 0.083 | 75.7 | 0.24 | 11 | 4 | 0.636 | 0.500 | 21.4 | 0.79 |
| | | 110 | 157 | 97 | 0.380 | 0.102 | 73.1 | 0.27 | 150 | 95 | 0.367 | 0.086 | 76.5 | 0.23 | 7 | 4 | 0.500 | 0.394 | 21.1 | 0.79 |
| H* | 3 | 117 | 179 | 115 | 0.356 | 0.117 | 67.0 | 0.33 | 168 | 111 | 0.339 | 0.102 | 70.0 | 0.30 | 11 | 4 | 0.606 | 0.374 | 38.2 | 0.62 |
| | | 124 | 175 | 114 | 0.349 | 0.105 | 69.9 | 0.30 | 161 | 109 | 0.321 | 0.075 | 76.5 | 0.23 | 14 | 5 | 0.667 | 0.459 | 31.2 | 0.69 |

[†] Phases: A, acclimation; L, low organic loading; H, high organic loading; H*, shift from high to low organic loading. [‡] Number of independent replicates. [§] Time (days).
[¶] Null model parameters: $\boldsymbol{\gamma_{obs}}$, observed gamma diversity; $\overline{\boldsymbol{\alpha_{obs}}}$, mean observed alpha diversity; $\boldsymbol{\beta_{obs}}$, observed beta diversity; $\boldsymbol{\beta_{obs}}$, observed beta diversity; DS, deterministic strength; $\overline{\boldsymbol{\beta_{exp}}}:\boldsymbol{\beta_{obs}}$, expected (mean) to observed beta diversity ratio.

**Table S4.** Parameter output from null model analysis on the genus-level dataset (metagenomics sequencing).

| **P**[†] | ***n***[‡] | **d**[§] | **All genera** (100% reads)[¶] | | | | | | **Common genera** (up to 99% acc. reads) | | | | | | **Rare genera** (less than 1% acc. reads) | | | | | |
|---|---|---|---|---|---|---|---|---|---|---|---|---|---|---|---|---|---|---|---|---|
| | | | $\boldsymbol{\gamma_{obs}}$ | $\overline{\boldsymbol{\alpha_{obs}}}$ | $\boldsymbol{\beta_{obs}}$ | $\overline{\boldsymbol{\beta_{exp}}}$ | **DS** (%) | $\overline{\boldsymbol{\beta_{exp}}}:\boldsymbol{\beta_{obs}}$ | $\boldsymbol{\gamma_{obs}}$ | $\overline{\boldsymbol{\alpha_{obs}}}$ | $\boldsymbol{\beta_{obs}}$ | $\overline{\boldsymbol{\beta_{exp}}}$ | **DS** (%) | $\overline{\boldsymbol{\beta_{exp}}}:\boldsymbol{\beta_{obs}}$ | $\boldsymbol{\gamma_{obs}}$ | $\overline{\boldsymbol{\alpha_{obs}}}$ | $\boldsymbol{\beta_{obs}}$ | $\overline{\boldsymbol{\beta_{exp}}}$ | **DS** (%) | $\overline{\boldsymbol{\beta_{exp}}}:\boldsymbol{\beta_{obs}}$ |
| A | 4 | 1 | 946 | 852 | 0.100 | 0.031 | 68.8 | 0.31 | 541 | 523 | 0.034 | 0.010 | 72.0 | 0.28 | 405 | 329 | 0.188 | 0.060 | 67.9 | 0.32 |
| | | 47 | 849 | 759 | 0.107 | 0.057 | 46.4 | 0.54 | 550 | 543 | 0.014 | 0.006 | 53.0 | 0.47 | 299 | 216 | 0.278 | 0.153 | 44.9 | 0.55 |
| L | 4 | 56 | 831 | 744 | 0.105 | 0.049 | 53.0 | 0.47 | 550 | 541 | 0.017 | 0.007 | 61.9 | 0.38 | 281 | 204 | 0.276 | 0.133 | 51.6 | 0.48 |
| | | 75 | 865 | 769 | 0.111 | 0.070 | 37.0 | 0.63 | 557 | 546 | 0.020 | 0.010 | 47.8 | 0.52 | 308 | 223 | 0.275 | 0.177 | 35.5 | 0.65 |
| | | 96 | 915 | 815 | 0.110 | 0.077 | 29.8 | 0.70 | 565 | 554 | 0.020 | 0.012 | 42.3 | 0.58 | 353 | 262 | 0.258 | 0.182 | 29.5 | 0.71 |
| | | 110 | 849 | 748 | 0.119 | 0.070 | 41.0 | 0.59 | 563 | 547 | 0.028 | 0.013 | 53.8 | 0.46 | 286 | 201 | 0.299 | 0.185 | 38.1 | 0.62 |
| | | 124 | 870 | 764 | 0.122 | 0.087 | 28.9 | 0.71 | 558 | 546 | 0.022 | 0.015 | 30.0 | 0.70 | 312 | 218 | 0.302 | 0.216 | 28.6 | 0.71 |
| H | 3 | 56 | 820 | 743 | 0.094 | 0.037 | 60.6 | 0.39 | 550 | 544 | 0.012 | 0.004 | 63.9 | 0.36 | 270 | 199 | 0.262 | 0.108 | 58.8 | 0.41 |
| | | 75 | 822 | 748 | 0.090 | 0.030 | 66.7 | 0.33 | 551 | 541 | 0.018 | 0.005 | 72.9 | 0.27 | 271 | 207 | 0.237 | 0.083 | 65.0 | 0.35 |
| | | 96 | 845 | 762 | 0.098 | 0.043 | 56.2 | 0.44 | 558 | 548 | 0.019 | 0.006 | 66.9 | 0.33 | 287 | 214 | 0.253 | 0.118 | 53.2 | 0.47 |
| | | 110 | 890 | 783 | 0.120 | 0.071 | 41.3 | 0.59 | 563 | 547 | 0.028 | 0.011 | 58.9 | 0.41 | 288 | 206 | 0.286 | 0.154 | 46.1 | 0.54 |
| H* | 3 | 124 | 890 | 783 | 0.120 | 0.071 | 41.3 | 0.59 | 561 | 546 | 0.027 | 0.015 | 46.3 | 0.54 | 329 | 237 | 0.279 | 0.168 | 39.8 | 0.60 |

[†] Phases: A, acclimation; L, low organic loading; H, high organic loading; H[*], shift from high to low organic loading.
[‡] Number of independent replicates
[§] Time (days)
[¶] Null model parameters: $\boldsymbol{\gamma_{obs}}$, observed gamma diversity; $\overline{\boldsymbol{\alpha_{obs}}}$, mean observed alpha diversity; $\boldsymbol{\beta_{obs}}$, observed beta diversity; $\boldsymbol{\beta_{obs}}$, observed beta diversity; DS, deterministic strength; $\overline{\boldsymbol{\beta_{exp}}}:\boldsymbol{\beta_{obs}}$, expected (mean) to observed beta diversity ratio.

**Table S5.** Parameter output from null model analysis on the IP2G gene-level dataset (metagenomics sequencing).

| P[†] | $n$[‡] | d[§] | **All IP2G genes** (100% reads)[¶] | | | | | | **Common IP2G genes** (up to 99% acc. reads) | | | | | | **Rare IP2G genes** (less than 1% acc. reads) | | | | | |
|---|---|---|---|---|---|---|---|---|---|---|---|---|---|---|---|---|---|---|---|---|
| | | | $\gamma_{obs}$ | $\overline{\alpha_{obs}}$ | $\beta_{obs}$ | $\overline{\beta_{exp}}$ | **DS** (%) | $\overline{\beta_{exp}}:\beta_{obs}$ | $\gamma_{obs}$ | $\overline{\alpha_{obs}}$ | $\beta_{obs}$ | $\overline{\beta_{exp}}$ | **DS** (%) | $\overline{\beta_{exp}}:\beta_{obs}$ | $\gamma_{obs}$ | $\overline{\alpha_{obs}}$ | $\beta_{obs}$ | $\overline{\beta_{exp}}$ | **DS** (%) | $\overline{\beta_{exp}}:\beta_{obs}$ |
| A | 4 | 1 | 3013 | 2634 | 0.126 | 0.157 | 24.4 | 1.24 | 1803 | 1791 | 0.007 | 0.013 | 105.8 | 2.06 | 1210 | 843 | 0.304 | 0.370 | 21.8 | 1.22 |
| | | 47 | 2563 | 2332 | 0.090 | 0.122 | 34.7 | 1.35 | 1801 | 1795 | 0.003 | 0.008 | 147.3 | 2.47 | 762 | 537 | 0.296 | 0.390 | 31.8 | 1.32 |
| L | 4 | 56 | 2518 | 2346 | 0.069 | 0.102 | 48.6 | 1.49 | 1800 | 1796 | 0.002 | 0.006 | 205.9 | 3.06 | 718 | 549 | 0.235 | 0.341 | 45.3 | 1.45 |
| | | 75 | 2440 | 2245 | 0.080 | 0.105 | 31.5 | 1.31 | 1805 | 1798 | 0.004 | 0.009 | 133.4 | 2.33 | 635 | 448 | 0.295 | 0.377 | 27.6 | 1.28 |
| | | 96 | 2388 | 2212 | 0.074 | 0.098 | 33.8 | 1.34 | 1803 | 1798 | 0.003 | 0.007 | 162.5 | 2.62 | 585 | 414 | 0.292 | 0.380 | 30.1 | 1.30 |
| | | 110 | 2395 | 2205 | 0.080 | 0.104 | 30.5 | 1.31 | 1804 | 1796 | 0.005 | 0.010 | 114.9 | 2.15 | 591 | 409 | 0.308 | 0.390 | 26.6 | 1.27 |
| | | 124 | 2399 | 2215 | 0.077 | 0.101 | 31.2 | 1.31 | 1804 | 1796 | 0.005 | 0.009 | 99.6 | 2.00 | 595 | 419 | 0.296 | 0.378 | 27.9 | 1.28 |
| H | 3 | 56 | 2481 | 2351 | 0.053 | 0.083 | 57.5 | 1.57 | 1798 | 1795 | 0.001 | 0.005 | 268.1 | 3.68 | 683 | 555 | 0.187 | 0.288 | 54.0 | 1.54 |
| | | 75 | 2422 | 2254 | 0.070 | 0.096 | 38.0 | 1.38 | 1798 | 1788 | 0.005 | 0.012 | 129.5 | 2.30 | 624 | 465 | 0.254 | 0.337 | 32.5 | 1.33 |
| | | 96 | 2344 | 2211 | 0.057 | 0.082 | 43.7 | 1.44 | 1800 | 1793 | 0.004 | 0.009 | 131.0 | 2.31 | 544 | 417 | 0.233 | 0.324 | 39.1 | 1.39 |
| | | 110 | 2343 | 2202 | 0.060 | 0.079 | 32.3 | 1.32 | 1799 | 1793 | 0.003 | 0.007 | 111.4 | 2.11 | 544 | 409 | 0.248 | 0.319 | 29.0 | 1.29 |
| H* | 3 | 124 | 2434 | 2255 | 0.074 | 0.094 | 27.8 | 1.28 | 1798 | 1792 | 0.003 | 0.006 | 103.2 | 2.03 | 636 | 463 | 0.273 | 0.342 | 25.5 | 1.25 |

[†] Phases: A, acclimation; L, low organic loading; H, high organic loading; H*, shift from high to low organic loading.
[‡] Number of independent replicates
[§] Time (days)
[¶] Null model parameters: $\gamma_{obs}$, observed gamma diversity; $\overline{\alpha_{obs}}$, mean observed alpha diversity; $\beta_{obs}$, observed beta diversity; $\beta_{obs}$, observed beta diversity; DS, deterministic strength; $\overline{\beta_{exp}}:\beta_{obs}$, expected (mean) to observed beta diversity ratio.
